# Construction of epithelial-mesenchymal transition related miRNAs signatures as prognostic biomarkers in gastric cancer patients

**DOI:** 10.1101/2022.09.02.506322

**Authors:** Jun Xiao, Fan Zhang, Wenju Liu, Weidong Zang

**Author notes:** Equally contributed (Jun Xiao, Fan Zhang). **Correspondence to**: Wei-Dong Zang, Department of Gastrointestinal Surgery, Fujian Medical University Cancer Hospital & Fujian Cancer Hospital, Fuzhou, Fujian, China.

## Abstract

**Aim:** To identify the potential post-healing EMT related miRNAs associated with lymph node metastatic gastric cancer (LNMGC).

**Methods:** Both RNA expression and clinical medical data were obtained from the TCGA dataset. We performed differential expression and normalization analysis of miRNAs. Cox linear regression model confirmed the differentially expressed miRNAs (DEmiRNAs) and clinical medical parameters related to overall survival (OS). The role of target genes of DEmiRNAs was determined according to the role enrichment analysis.

**Results:** We obtained a total of 7531 DEmRNAs and 267 DEmRNAs, of which 185 DEmRNAs were down-regulated and 82 DEmRNAs were up-regulated. We randomly divided the LMNGC cases (n=291) into a training group (n=207) and a test group (n=84). The results showed that a total of 103, 11, 13 and 83 overlapping genes were associated with hsa-mir-141-3p, hsa-mir-4664-3p, hsa-mir-125b-5p and hsa-mir-7-5p, respectively. Kaplan-Meier determined that these four miRNAs can effectively distinguish high-risk and low-risk groups, and have a good indicator role (all p<0.05). Multivariate cox regression analysis also showed that EMT-related miRNA predictive model and lymph node metastasis were both prognostic risk factors (all p<0.05). The ROC curve showed that this feature had high accuracy (AUC>0.7, p<0.05). In addition, KEGG analysis showed that EMT-related pathways were mainly enriched in HIF-1 signaling pathway and focal adhesion.

**Conclusions:** Our study demonstrated that EMT-related miRNAs could serve as independent prognostic markers in pN_1-3_ GC patients.

## Introduction

In the statistical analysis of gastric cancer (GC) in the world in 2020, its incidence rate and mortality ranked sixth and third respectively[1]. The prognosis time of GC is still very low, mainly because GC patients are usually diagnosed as intermediate and advanced stage[2]. According to the references, about 39% of GC patients were diagnosed with tumor metastasis, and the survival rate of this part was generally very low[3]. Internationally, TNM staging is used to evaluate the prognosis of malignant tumors[4]. However, due to individual heterogeneity, the prognosis of GC patients varies greatly even within the same TNM stage[5]. Therefore, we try to adopt new strategies to improve survival analysis and prediction to further guide the individualized treatment of GC.

Epithelial mesenchymal transition (EMT) is involved in the processes related to the invasion, metastasis, drug resistance and prognosis of malignant tumors[6]. Due to the diversity of individuals and the difference of tumor microenvironment, EMT has been proved to have healing effect in GC, which leads to lack of relevant genetic healing entity models in specific clinical applications[7]. MicroRNAs (miRNAs) are a group of non numbered polypeptide chain microRNAs (18-22 ribonucleotides) that inhibit gene expression based on 3’-UTR fusion with target mRNAs[8, 9]. There is increasing evidence that various miRNAs play a vital role in EMT[10, 11]. Therefore, a better understanding of the efficacy of EMT-related miRNAs can help us further explore its uncertain diagnostic, healing and diagnostic value.

It has been emphasized that EMT is closely related to cancer metastasis, and this process is closely related to the regulation of the tumor microenvironment[10, 12, 13]. Some miRNAs have long been confirmed to participate in the EMT process, but the mechanism of how EMT related genetic genes and miRNAs interaction is still unclear[14]. This scientific research is committed to screening and evaluating important miRNAs in the EMT process in general, and further screening different biological transcription factors according to distinct genes.

## Materials and methods

### Source and collection of database

Original RNA transcriptase transcriptome sequencing (RNA-seq) data information system variables are loaded from transcriptase transcriptome sequencing in TCGA database. In a previous systematic study, 200 genes related to EMT were evaluated (Table S1). Clinical medical professional information of LNMGC cases, including gender, age, TNM stage and prognostic information. The mRNA expression files included 375 GC cases and 32 controls, while the miRNA expression profile included 446 GC cases and 45 controls. Subsequently, we downloaded all mature miRNA coding sequences from miRBase (http://www.mirbase.org/). Using the Perl language, we merged transcriptome and clinical information, which allowed us to obtain a merged file of miRNAs and GC clinical and prognostic information.

### Assessing differential factors for mRNAs and miRNAs

We extracted EMT-related candidate genes (h.all.v7.1.symbols) from the GSEA database. Screening criteria were: false detection rate (FDR)<0.05 and 2-fold transformations (FC)>1. DEmRNAs and DEmirnas from LMNGC patients and normal controls were screened and normalized according to the edger package (Table S2). Additionally, we use the ggplot and pheatmap packages to make volcano plots and pheatmap plots.

### Establishment of EMT-related miRNA prediction model

We screened 267 EMT-related miRNAs and clinicopathological parameters of GC (Table S3), divided the training and validation sets into subgroups in a 7:3 ratio. Univariate and multivariate cox regression analysis were used to screened out the important EMT-related miRNAs in the training set. Based on the above miRNAs, we further constructed a predictive model. According to the risk value of the risk score, GC patients were subdivided into high-risk and low-risk groups. In addition, the ROC curve also verified the accuracy of the risk model. According to the predictive model risk score, patients were divided into high-risk set and low-risk set. The accuracy of the predictive model was verified by ROC curve.

### EMT-related miRNAs and target genes and potential roles

We identified miRNA target genes through online databases such as targetscan dataset, miRDB and mirtarbase. The screened EMT-related genes should cover at least 2 datasets, and a venn diagram should be drawn. In addition, we also demonstrated the regulatory relationship between EMT-related miRNAs and target genes by cytoscape. To reveal the mechanism of action of target genes in EMT process, we performed enrichment analysis by GO and KEGG.

### Statistical Analysis

All statistical processing was performed by R software (vision 4.2). Univariate and multivariate risk assessments were performed by cox regression models. OS was assessed in high-risk and low-risk patients by Kaplan-Meier and log-rank tests. Risk values were assessed with hazard ratios (HR) and 95% confidence intervals (95% CI). The Mann Whitney U test was used to compare two groups. P<0.05 was considered statistically significant.

## Results

### Identification of DEmiRNAs and DEmRNAs

Through differential analysis of the TCGA database, we obtained a total of 7531 DEmRNAs and 267 DEmRNAs. Among them, 3136 DEmRNAs were down-regulated and 4395 were up-regulated (Figure 1A/B). What is more, 82 DEmiRNAs were down-regulated and 185 were up-regulated (Figure 1C/D). We randomly divided the LMNGC cases (n=291) into a training group (n=207) and a test group (n=84). We then constructed a cox risk prediction model as the outcome variable. Univariate regression analysis showed that hsa-mir-139-5p, hsa-mir-141-3p, hsa-mir-143-5p, hsa-mir-328-3p, hsa-mir-96-5p, hsa-mir-100-5p, hsa-mir-4664-3p, hsa-mir-125b-5p, hsa-mir-7-5p and hsa-mir-504-5p were significantly associated with OS.

**Figure 1.**
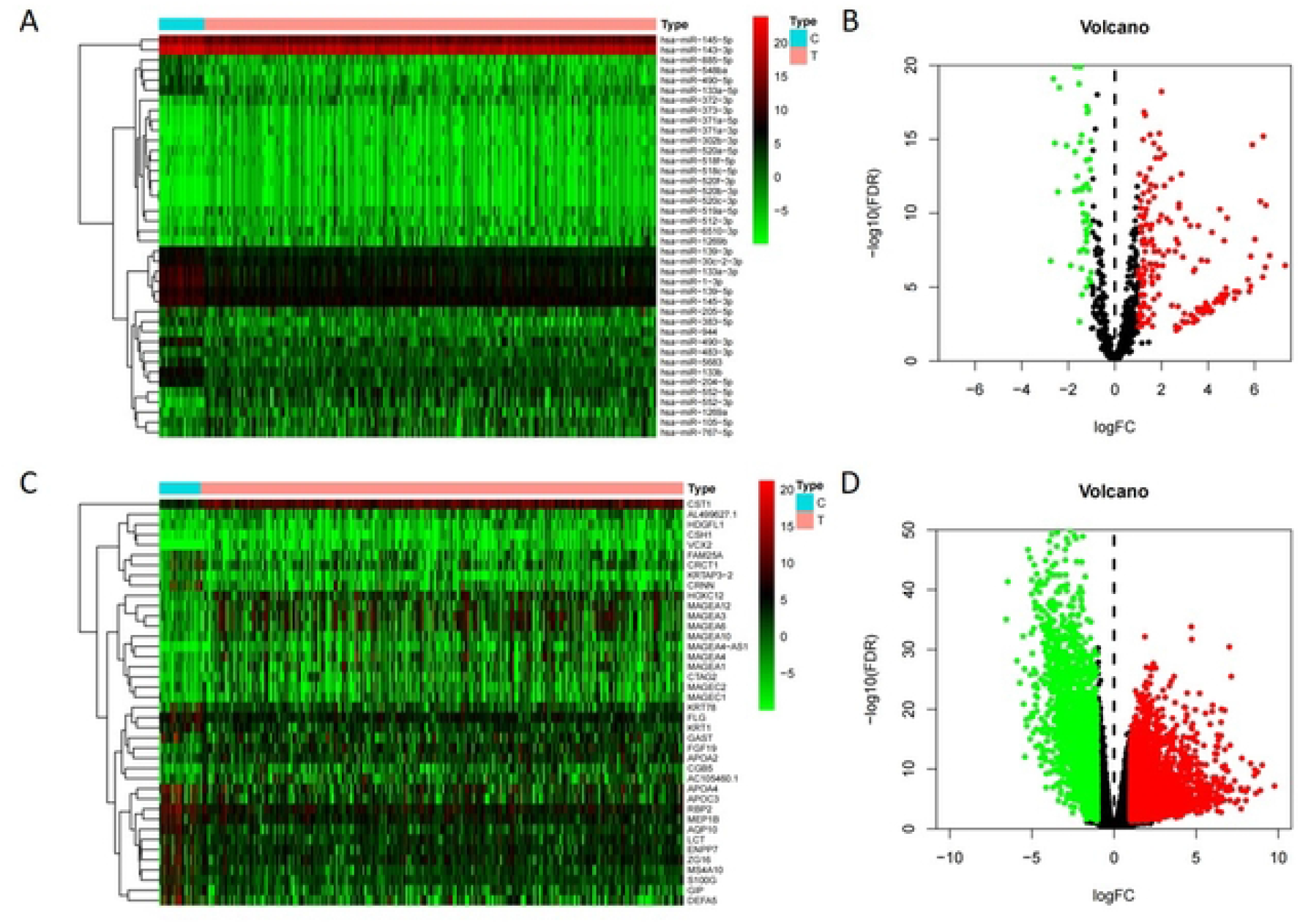
Heatmap and volcano plots of DEmRNAs and DEmiRNAs between LNMGC patients and normal controls. (A) and (C) Heatmap of top 40 DEmRNAs and DEmiRNAs, mRNA and miRNA that up-regulated are in red. mRNA and miRNA that down-regulated are in green. mRNA and miRNA that without any significant difference are in black. (B) and (D) Volcano plots of DEmRNAs and DEmiRNAs.

### Prediction of target DEmiRNAs

We combined network databases (targetscan, mirdb, mirtarbase) and EMT-related genes for analysis, visualized the regulatory network of miRNAs and target genes (Fig. 2). It was obtained the intersection of miRNAs and visualized by venn diagram (Fig. 3). The results showed that a total of 103, 11, 13 and 83 overlapping genes were associated with hsa-mir-141-3p, hsa-mir-4664-3p, hsa-mir-125b-5p and hsa-mir-7-5p, respectively. In addition, 139 target genes were found to be differentially expressed, of which 108 genes were down-regulated and 31 genes were up-regulated. Differential miRNAs were further observed and their clinical value in prognosis was assessed (Fig. 4A/B). Kaplan-Meier determined that these four miRNAs can effectively distinguish high-risk and low-risk groups, and have a good indicator role (Figure 5).

**Figure 2.**
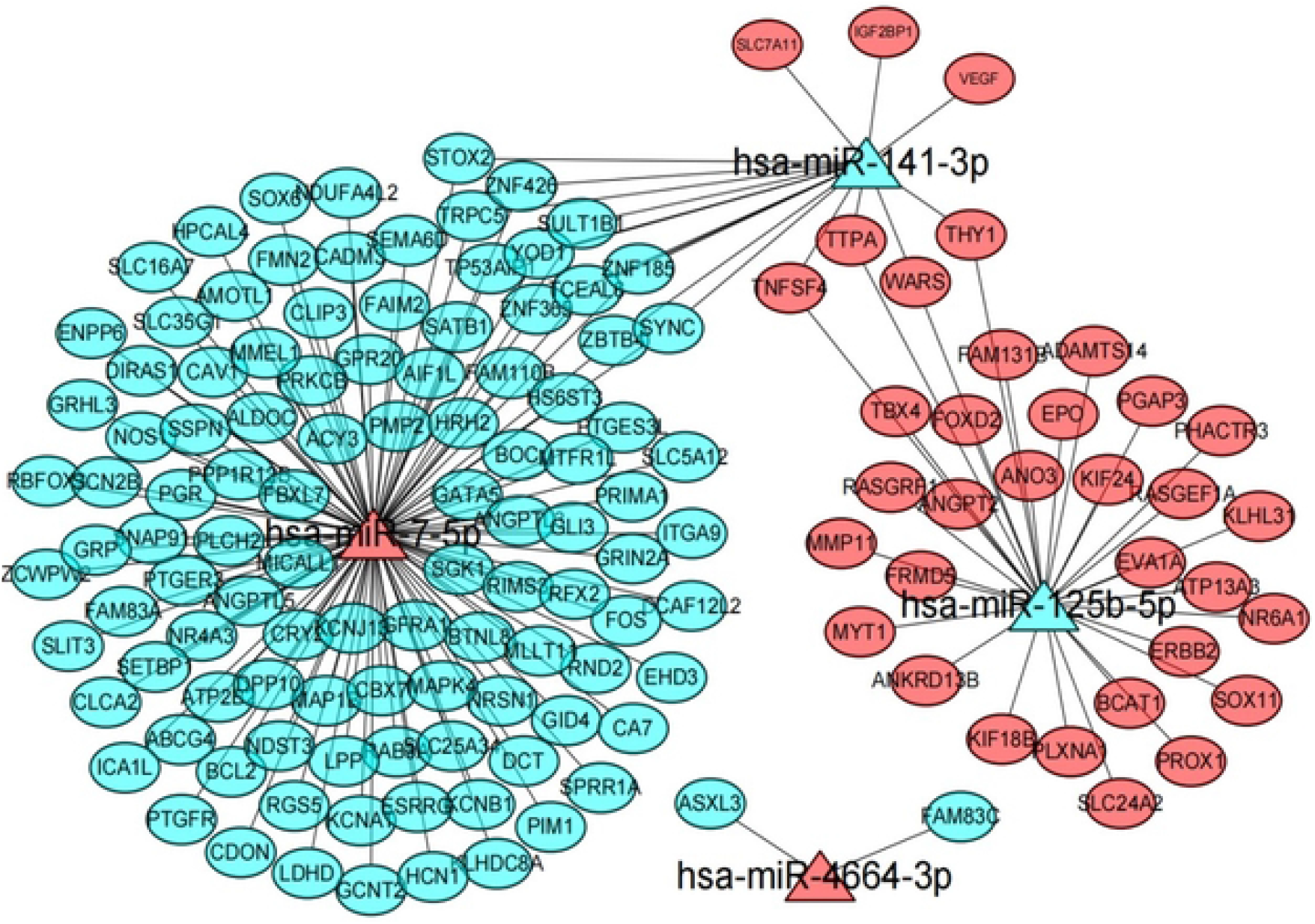
Interaction networks of the four EMT-related miRNAs and their target genes. The ellipse represents target gene, triangle represents miRNA, and the link in black indicates a miRNA-target gene relationship.

**Figure 3.**
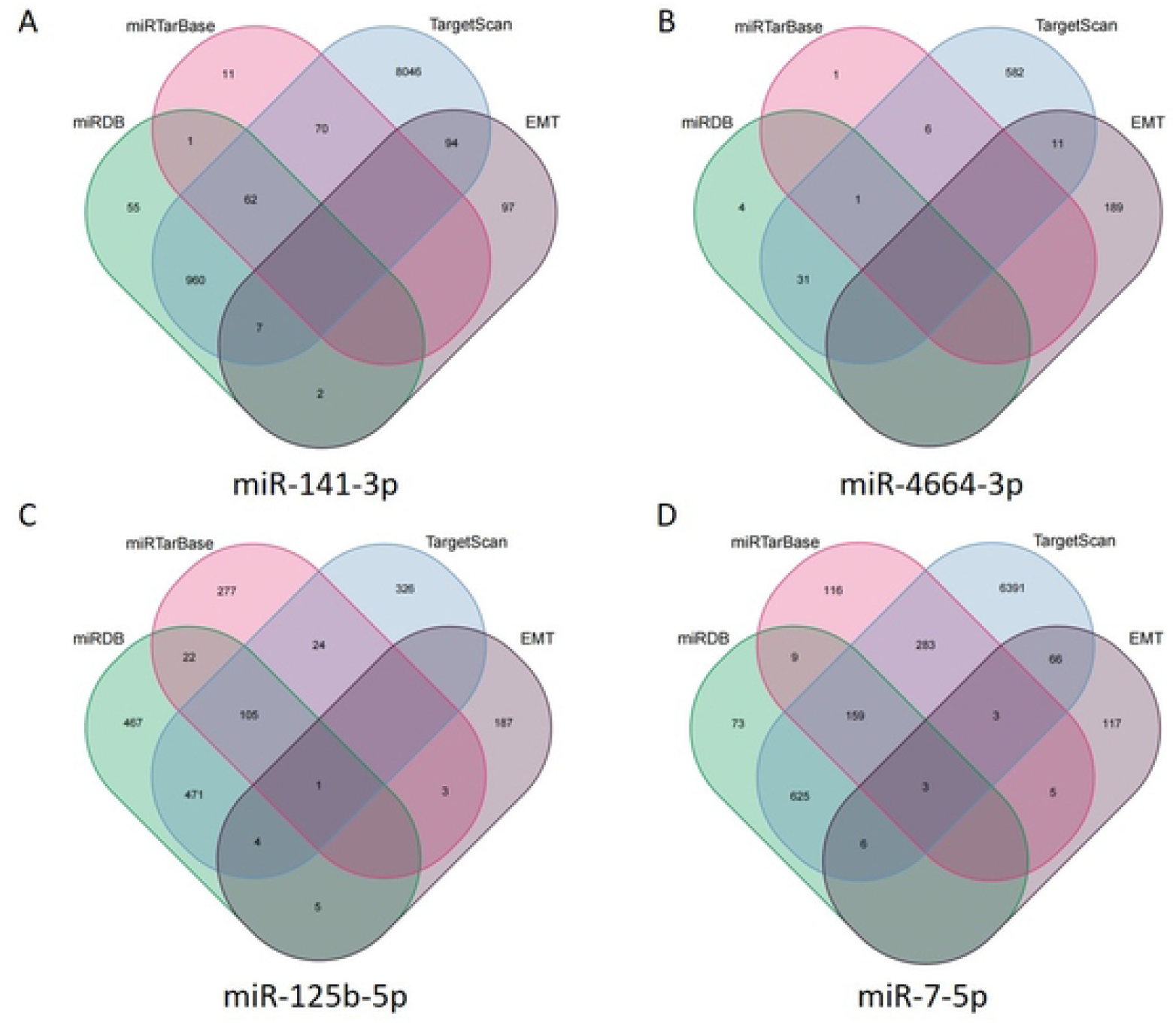
Venn diagram of the intersection of four EMT-related miRNAs. (A) hsa-miR-141-3p, (B) hsa-miR-4664-3p, (C) hsa-miR-125b-5p, (D) hsa-miR-7-5p.

**Figure 4.**
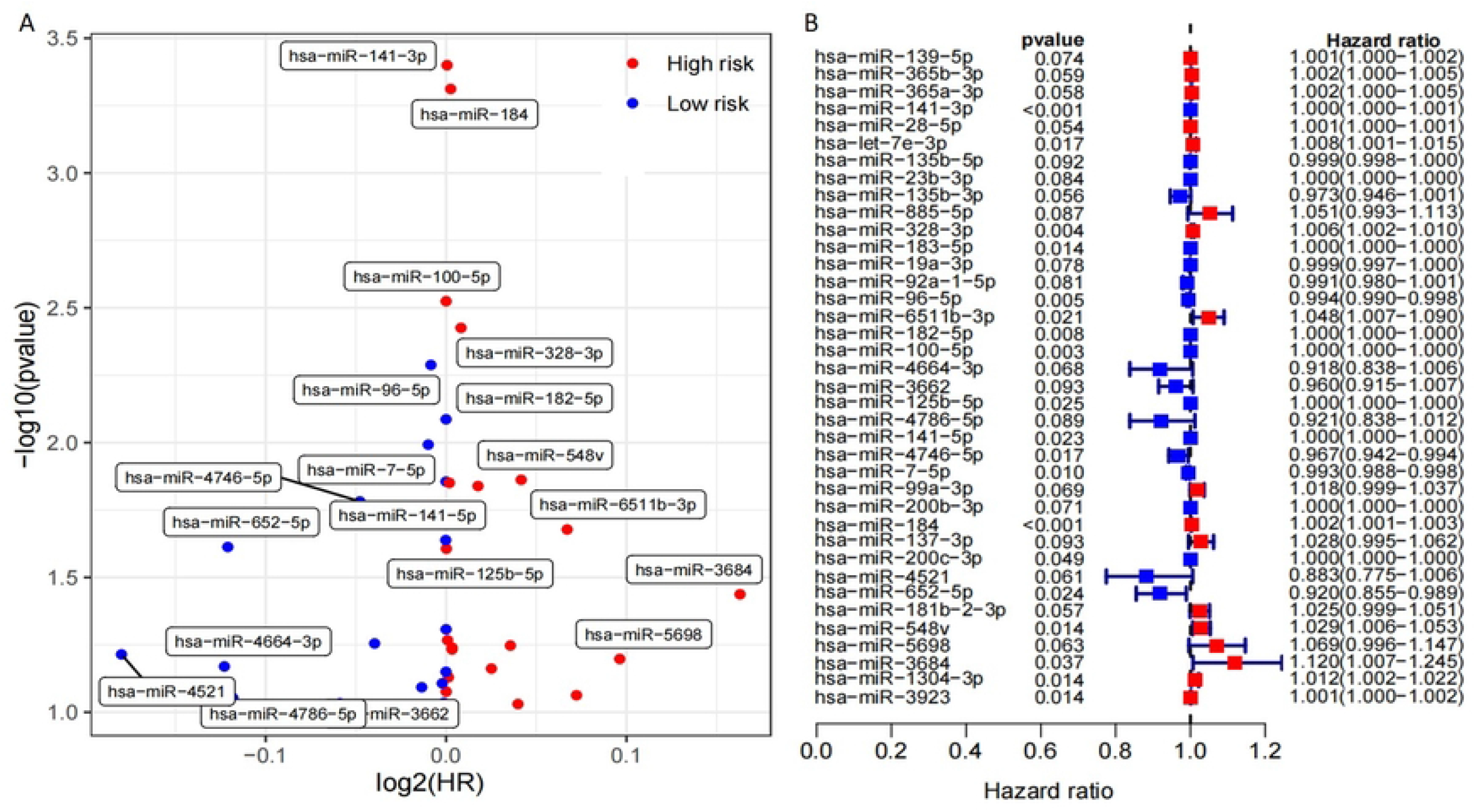
Screening of EMT-related differential miRNAs. A volcano plot of expression change comparing the tumor and normal samples, with the downregulated and upregulated miRNAs. (B) Univariate analysis for the TCGA cohort.

**Figure 5.**
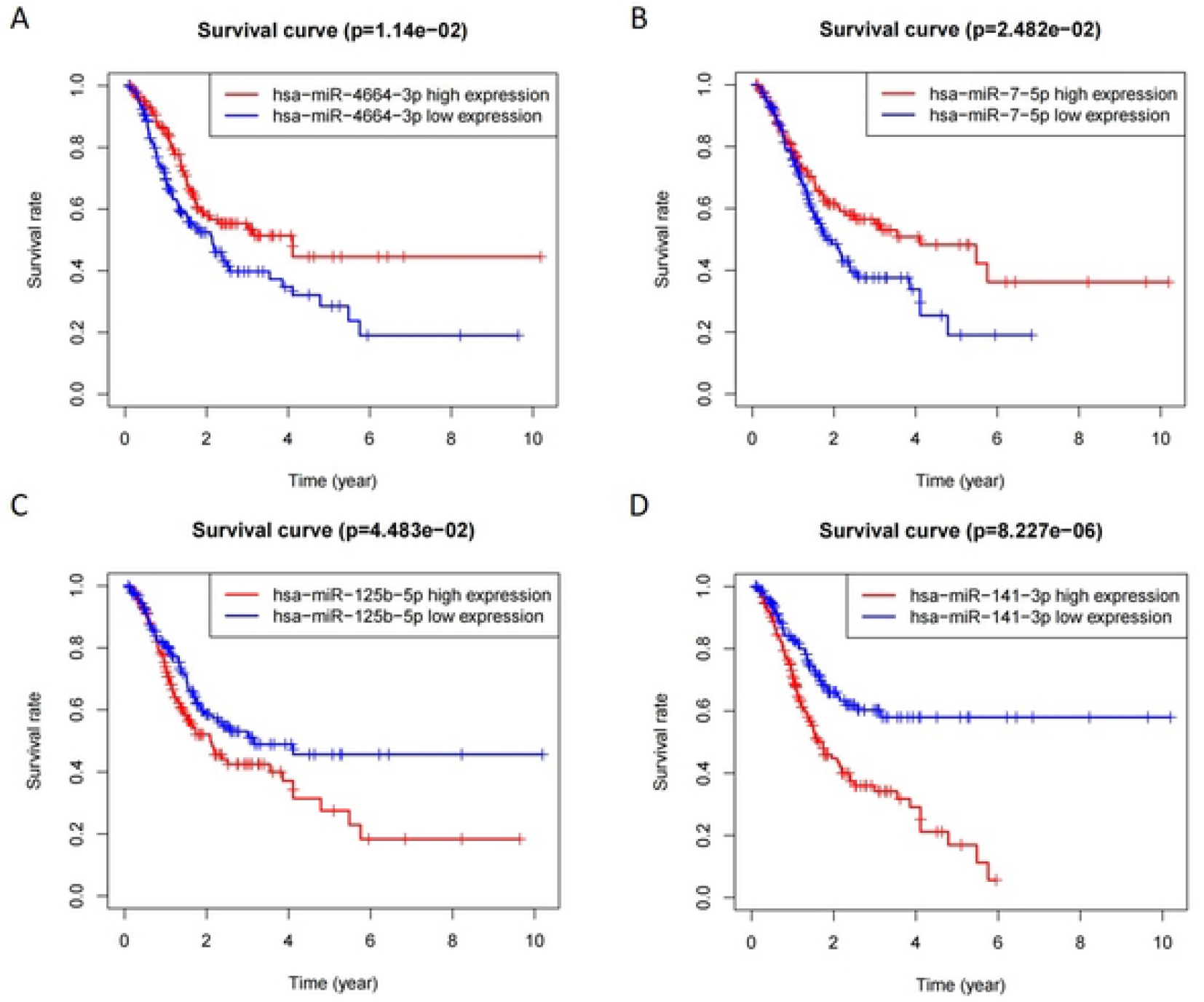
Four EMT-related miRNAs associated with OS. High- and low-expression groups were subdivided according to the median expression of each miRNA. (A) hsa-miR-141-3p, (B) hsa-miR-4664-3p, (C) hsamiR-125b-5p, (D) hsa-miR-7-5p. P-value was assessed with log-rank test.

### Construction of EMT-related miRNA prognostic model

We combined analysis of EMT-related miRNA predictors and other clinical characteristics, and it was divided into the high-risk group and low-risk group according to different risk scores (Figure 6A). The setting parameters of the EMT-related miRNAs model are: risk score= 0.533 × (hsa-mir-141-3p)-0.188 × (hsa-mir-7-5p)-0.183 × (hsa-mir-4664-3p)-0.188× (hsa-mir-125b-5p).

**Figure 6.**
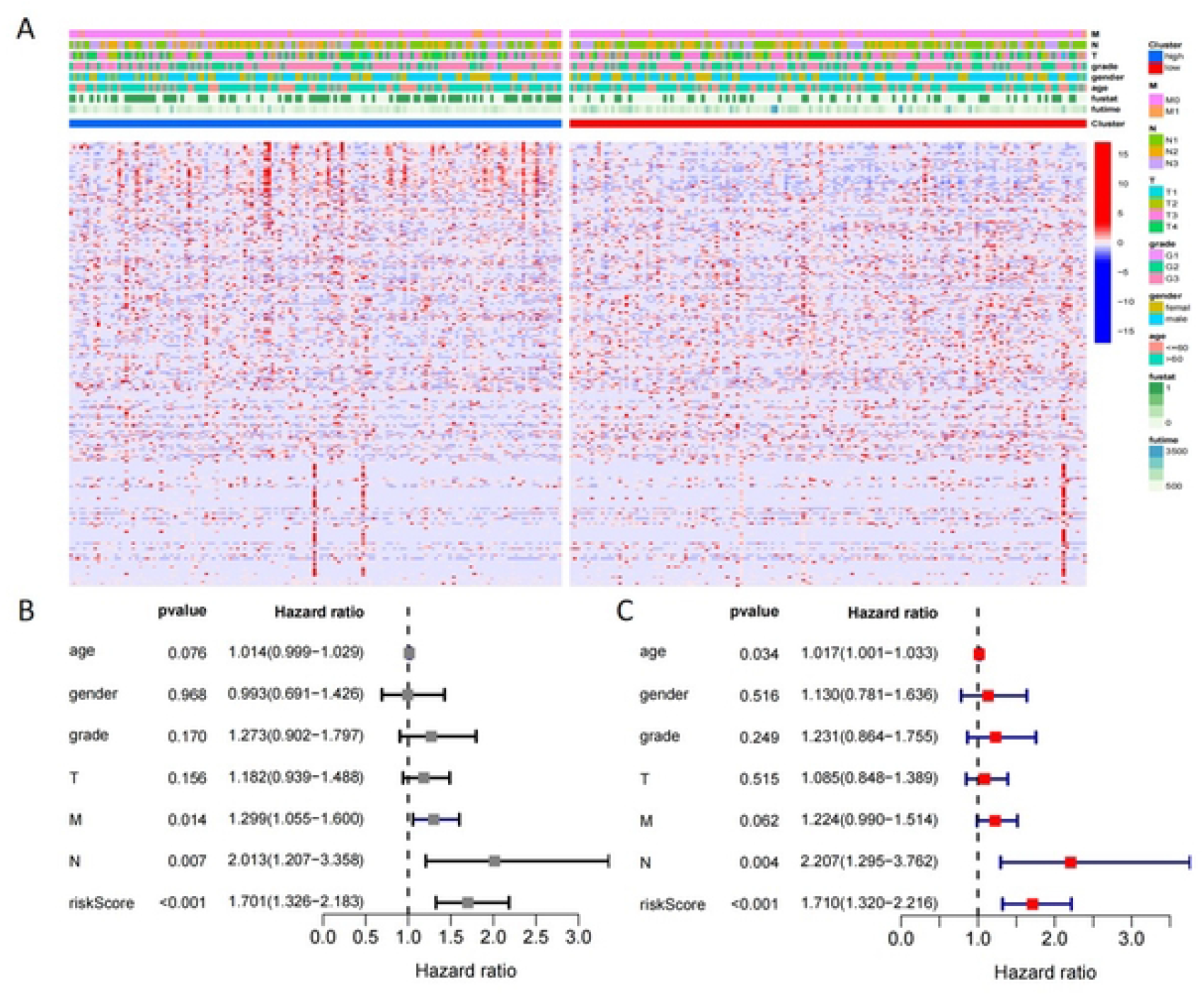
Construction of a prognostic model for EMT-related miRNAs. (A) Heatmap of clinicopathological characteristics of the low- and high-risk patients. (B) Univariate analysis for the TCGA cohort. (C) Multivariate analysis for the TCGA cohort. HR: hazard ratio; CI: confidence interval.

Univariate Cox multiple regression analysis indicated that both EMT-related miRNA predictive model (Fig. 6B; HR=1.701, p<0.001) and lymph node metastasis were prognostic risk factors (Fig. 6B; HR=2.013, p=0.007). Meanwhile, multivariate cox regression analysis also showed that EMT-related miRNA predictive model and lymph node metastasis were both prognostic risk factors (Fig. 6C; all p<0.05).

### Evaluation of the validity of a prognostic model

By Kaplan-Meier survival analysis, we evaluated all LMNGC patients (Fig. 7A, 13.77% vs 56.09%, P<0.05), training group (Fig. 7B, 11.89% vs 53.84%) and test group (Fig. 7C, 18.97 % vs 60.10%), respectively. The results showed that the prognosis of the high-risk group was significantly lower than that of the low-risk group in the above three data sets (all p<0.05). In addition, we also plotted the distribution of survival for all patients (Fig. 7D), the training group (Fig. 7E), and the test group (Fig. 7F). We evaluated the effectiveness of the model by the survivalROC package, and the AUC areas of all cases (Fig. 7G, 0.755), training set (Fig. 7H, 0.750) and test set (Fig. 7I, 0.778) were all above 0.7, indicating that the model has excellent predictive ability.

**Figure 7.**
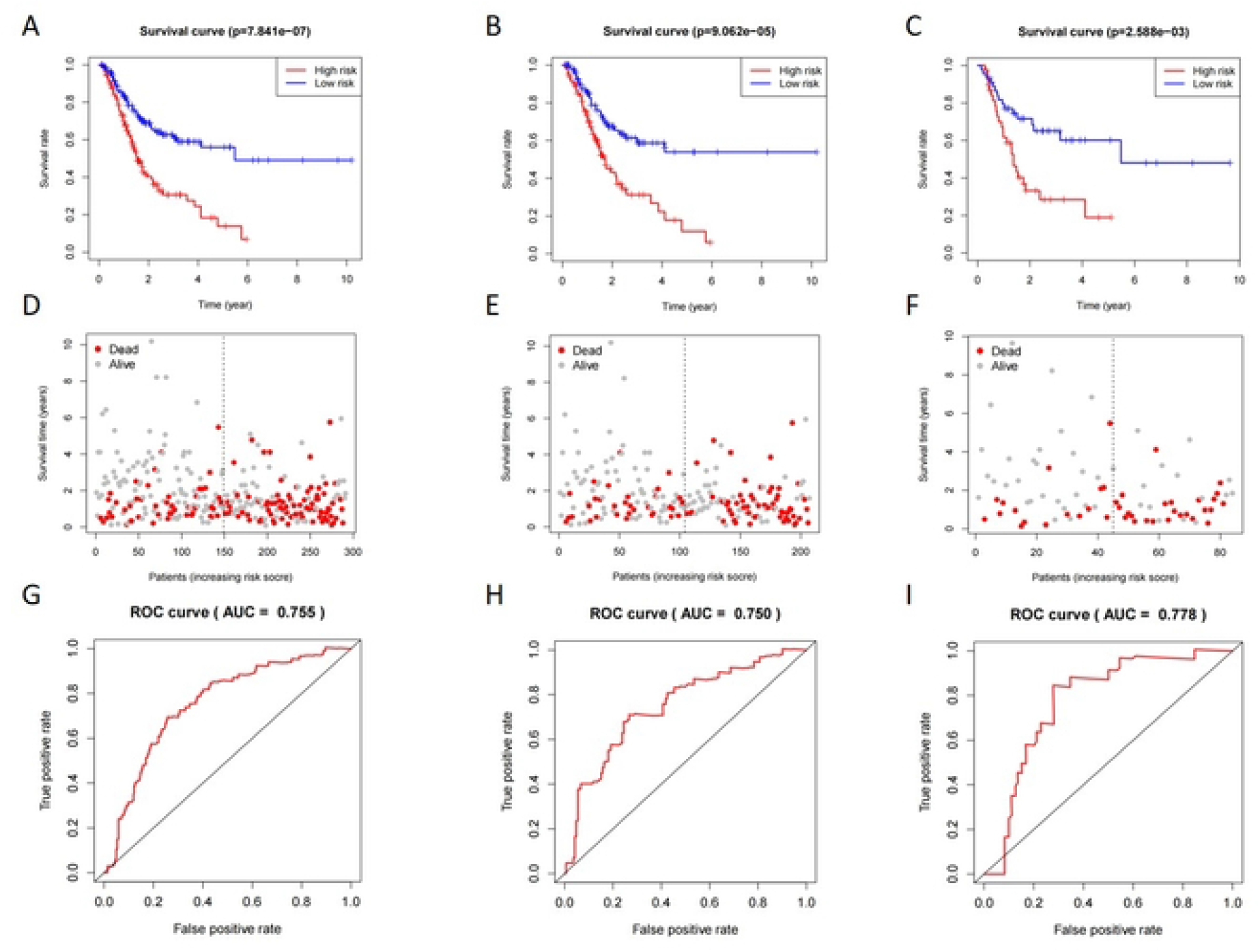
Evaluation and validation of the Four EMT-related miRNAs signature. Kaplan-Meier curves of the (A) all cases, (B) training set, (C) testing set; Survival status Plot of the (D) all cases, (E) training set, (F) testing set; The AUC of the 5-year dependent ROC curve in the (G) all cases, (H) training set, (I) testing set (F). P-value was assessed with log-rank test.

### Gene ontology and KEGG pathway analysis

According to the clusterprofiler package, we obtained GO and KEGG enrichment of all target genes participating in lymph node metastasis. The GO analysis results show that 483 GO annotations are related to the development trend of EMT. The dot chart shows the top terms of the biological process, cellular component and molecular function of all microbial strains (Figures 8A/B/C).

**Figure 8.**
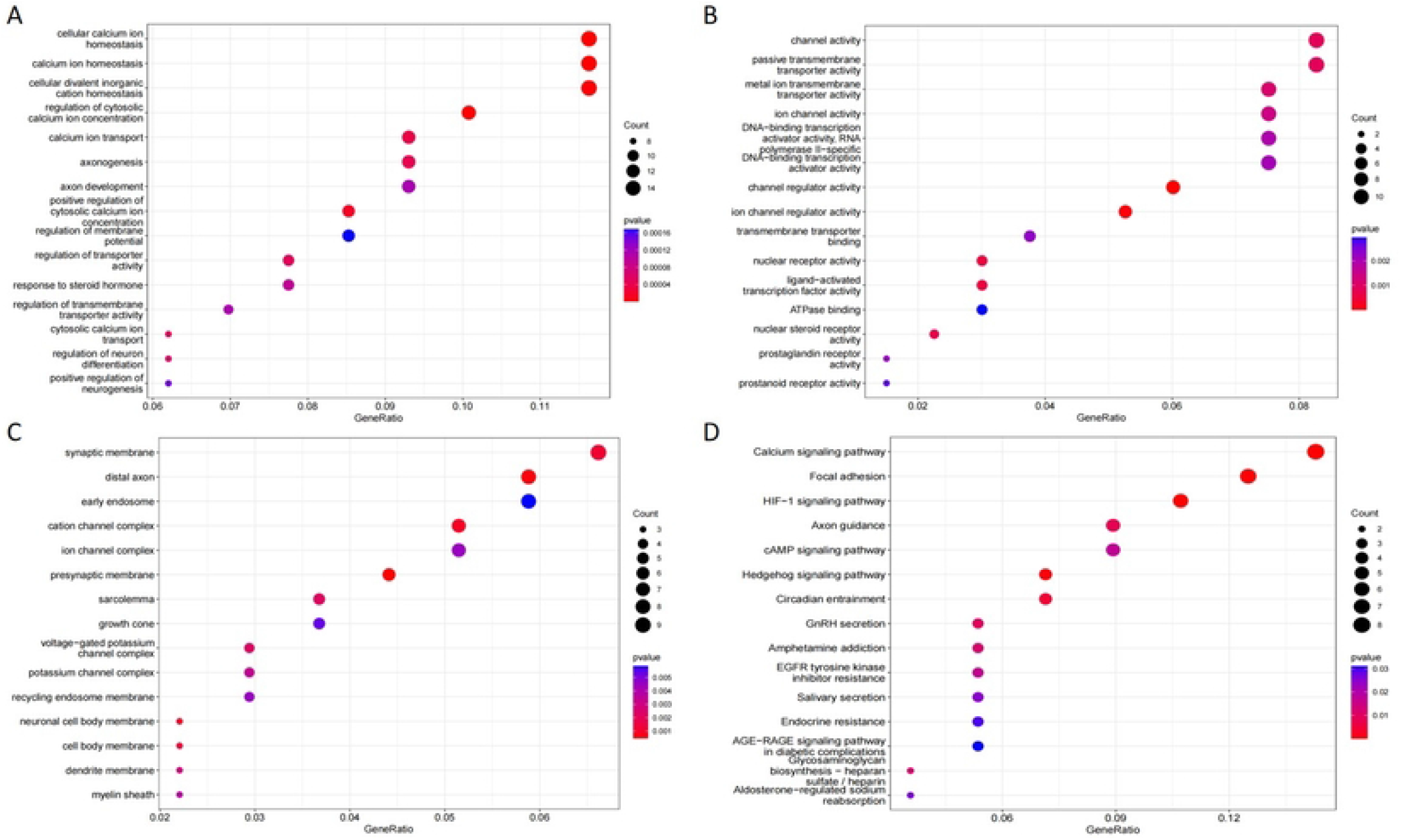
GO and KEGG pathway analysis of target genes. BP(A), MF(B), and CC(C) involved in molecular characteristics of target genes, (D)Top 19 signaling pathways of KEGG.

Among these three categories, EMT-related target genes are mainly enriched in BP genes, including include cellular calcium ion homeostasis, calcium ion homeostasis, and regulation of cytosolic calcium ion concentration. For CC section analysis showed that the presynaptic membrane, distal axon, and cation channel complex. In the MF section, channel regulator activity, ion channel regulator activity, and nuclear receptor activity are the first three technical terms for overall target gene enrichment.

In addition, KEGG analysis results showed that 19 GC related pathways were mainly enriched in HIF-1 data signal pathway, calcium data signal pathway and adhesive plaque (Figures 8D). In addition, we also brought “pathway gene Internet” (Figures 9a) and “pathway pathway Internet” (Figures 9b) to confirm the relationship between KEGG pathway and target genes.

**Figure 9.**
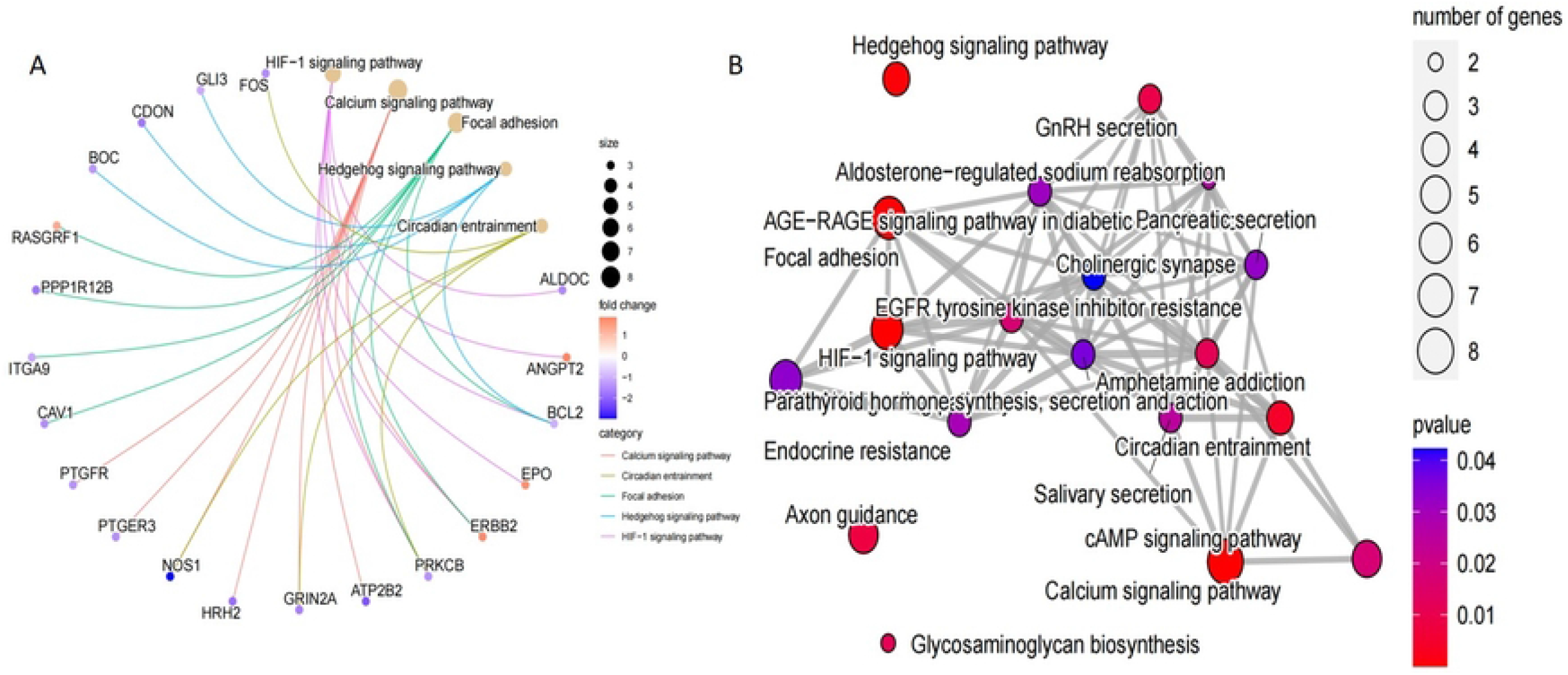
Enrichment analysis of genes to pathway network. (A) pathway-gene network, (B) pathwaypathway network.

## Discussion

EMT is a complex biological process, and the regulatory network consists of multiple miRNAs and target genes, and its role in predicting GC patients has received increasing attention[15]. Many studies have reported that the EMT process is related to the lymph node metastasis process of malignant tumors[16]. Many EMT-related studies usually focus on the study of a single gene, so the conclusions drawn are often inconsistent[17, 18]. This project is based on 200 common EMT-related genes to screen out miRNAs related to EMT genes. The integration of these miRNAs has important implications for understanding biological processes.

This study found that hsa-mir-141-3p, hsa-mir-7-5p, hsa-mir-4664-3p and hsa-mir-125b-5p had good discriminative ability in survival analysis (all, P<0.05). The novelty of this study is that after obtaining the characteristics of EMT-related miRNAs, the predicted target genes are also obtained from the online database, which rides the results of this study more accurate. Multivariate regression analysis showed that EMT-related miRNA prognostic model and lymph node metastasis were risk prognostic factors (all P<0.05). The AUC of the EMT-related miRNA prognostic model was the training set (0.778) and the validation set (0.750), respectively, indicating that the EMT-related miRNAs model has excellent predictive ability.

In a recent study[19], it also had a miRNA prognostic model for GC, which included multiple miRNAs (miR-453-3p, let-7a-3p, miR-145-3p, miR-135b-5p, miR-204-5p, miR-149-5p and miR-195-5p). However, this article included not only patients with pN0, but also patients with lymph node metastasis, which may make the prediction power of this article insufficient. The inclusion of cases in this study excluded pN0 cases, which further ruled out confounding factors and made the model more targeted.

In this study, we screened out four DEmiRNAs and 131 target genes. The function and pathway enrichment of EMT-related target genes were further analyzed using GO and KEGG methods. The results showed that GO annotations of differentially target genes were mainly related to cellular calcium homeostasis, calcium homeostasis, regulation of cytosolic calcium concentration, presynaptic membrane, distal axon, cation channel complexes, channel regulator activity, ion Channel modulator activity and nuclear receptor activity. KEGG analysis mainly enriched the target genes of HIF-1 signaling pathway, calcium signaling pathway and focal adhesion. Abnormal signaling pathways play a vital role in tumorigenesis, and increasing evidence suggests that HIF-1 signaling pathway and focal adhesion are involved in the development of GC.

EMT is a multifaceted microbial process involving multiple pathways[20, 21]. KEGG enrichment analysis in our study suggested that the interaction between the HIF-1 signaling pathway and focal adhesion may be two key features of significantly EMT-related pathways. The literature reports that focal adhesions play a major role in altering cell morphology and motility and regulating cell proliferation and differentiation[22–24]. HIF-1 signaling pathway participates in the process of angiogenesis in cancer tissue by regulating the expression of downstream angiogenesis factors, which promotes tumor cell proliferation, and this process is closely linked to the HIF-1/VEGF signaling pathway[25].

There are still some limitations to our study. First, due to the lack of certification of blood samples, the number of certified samples must be further expanded in subsequent stages. Second, we still do not know whether the differential expression of miRNAs is generated by the tumor metastasis process or by microenvironmental changes made by the host to the tumor metastasis process. Therefore, the roles of these four miRNAs must be further investigated.

## Conclusion

We build a prediction model of EMT-related miRNAs, and it can individually predict the prognosis of GC patients with lymph node metastasis. In addition, a large cohort study is still required to further validate the validity of the model before it can be gradually applied to clinical practice.

**Table 1.** Clinical characteristics of LNMGC patients.

| Covariates |  | Case | Percentage (%) |
| --- | --- | --- | --- |
| Age(years) |  |  |  |
|  | <60 | 94 | 32.30 |
|  | ≥ 60 | 197 | 67.70 |
| Gender |  |  |  |
|  | Female | 100 | 34.36 |
|  | Male | 191 | 65.64 |
| Grade |  |  |  |
|  | G1 | 5 | 1.72 |
|  | G2 | 98 | 33.68 |
|  | G3 | 188 | 64.60 |
| Lymph node status |  |  |  |
|  | N1 | 114 | 39.18 |
|  | N2 | 87 | 29.90 |
|  | N3 | 90 | 30.93 |
| T stage |  |  |  |
|  | T1 | 6 | 2.06 |
|  | T2 | 53 | 18.21 |
|  | T3 | 138 | 47.42 |
|  | T4 | 94 | 32.30 |
| TNM stage |  |  |  |
|  | I | 4 | 1.37 |
|  | II | 99 | 34.02 |
|  | III | 160 | 54.98 |
|  | IV | 28 | 9.62 |

LNMGC: lymph node metastatic gastric cancer

**Table 2.** DEmiRNAs involved in prognosis associated model of LNMGC.

| uniCox |  |  |  |  | multiCox |  |  |  |  |
| --- | --- | --- | --- | --- | --- | --- | --- | --- | --- |
| id | HR | HR.95L | HR.95H | pvalue | id | HR | HR.95L | HR.95H | pvalue |
| age | 1.014 | 0.999 | 1.029 | 0.076 | age | 1.017 | 1.001 | 1.033 | 0.034 |
| gender | 0.993 | 0.691 | 1.426 | 0.968 | gender | 1.130 | 0.781 | 1.636 | 0.516 |
| grade | 1.273 | 0.902 | 1.797 | 0.170 | grade | 1.231 | 0.864 | 1.755 | 0.249 |
| T stage | 1.182 | 0.939 | 1.488 | 0.156 | T stage | 1.085 | 0.848 | 1.389 | 0.515 |
| M stage | 1.299 | 1.055 | 1.600 | 0.014 | M stage | 1.224 | 0.990 | 1.514 | 0.062 |
| N stage | 2.013 | 1.207 | 3.358 | 0.007 | N stage | 2.207 | 1.295 | 3.762 | 0.004 |
| riskScore | 1.701 | 1.326 | 2.183 | 0.000 | riskScore | 1.710 | 1.320 | 2.216 | 0.000 |

Differentially expressed miRNAs, LNMGC: lymph node metastatic gastric cancer

## Abbreviations

GC: Gastric cancer,
LNMGC: lymph node metastasis of GC,
EMT: Epithelial mesenchymal transition,
miRNAs: MicroRNAs,
RNA-seq: RNA transcriptase transcriptome sequencing,
FDR: false discovery rate,
DEmRNAs: Differentially expressed mRNAs,
DEmiRNAs: Differentially expressed miRNAs,
AUC: area under roc curve.

## Data Availability

The data used to support the findings of this study are included within the supplementary information files.

## Acknowledgments

Not applicable.

## Author contributions

ZWD and XJ wrote the main manuscript, XJ and ZF performed the analyses. XJ, and LWJ collected data and prepared all figures and tables. XJ and ZWD conceived of and designed the study. All authors reviewed and approved the manuscript.

